# Understanding signaling and metabolic paths using semantified and harmonized information about biological interactions

**DOI:** 10.1101/2020.07.31.230599

**Authors:** Ryan A Miller, Martina Kutmon, Anwesha Bohler, Andra Waagmeester, Chris T Evelo, Egon L Willighagen

## Abstract

**Background:** To grasp the complexity of biological processes, the biological knowledge is often translated into schematic diagrams of biological pathways, such as signalling and metabolic pathways. These pathway diagrams describe relevant connections between biological entities and incorporate domain knowledge in a visual format that is easier for humans to interpret. It has already been established that these diagrams can be represented in machine readable formats, as done in KEGG, Reactome, and WikiPathways. However, while humans are good at interpreting the message of the creator of such a diagram, algorithms struggle when the diversity in drawing approaches increases. WikiPathways supports multiple drawing styles, and therefore needs to harmonize this to offer semantically enriched access via the Resource Description Framework format. Particularly challenging in the normalization of diagrams are the interactions between the biological entities, so that we can glean information about the connectivity of the entities represented. These interactions include information about the type of interaction (metabolic conversion, inhibition, etc.), the direction, and the participants. Availability of the interactions in a semantic and harmonized format enables searching the full network of biological interactions and integration with the linked data cloud.

**Results:** We here study how the graphically modelled biological knowledge in diagrams can be semantified and harmonized efficiently, and exemplify how the resulting data can be used to programmatically answer biological questions. We find that we can translate graphically modelled biological knowledge to a sufficient degree into a semantic model of biological knowledge and discuss some of the current limitations. Furthermore, we show how this interaction knowledge base can be used to answer specific biological questions.

**Conclusion:** This paper demonstrates that most of the graphical biological knowledge from WikiPathways is modelled in the semantic layer of WikiPathways with the semantic information intact and connectivity information preserved. The usability of the WikiPathways pathway and connectivity information has shown to be useful and has been integrated into other platforms. Being able to evaluate how biological elements affect each other is useful and allows, for example, the identification of up or downstream targets that will have a similar effect when modified.

## Background

Human cells contain around 20,000 protein-coding genes and that does not even include numerous non-coding genes [1] and the many encoded proteins. Furthermore, the Human Metabolome Database (HMDB) describes over 100,000 metabolites [2]. The number of interactions between biological entities is even higher. For example, cells also contain protein complexes, of which over 600 known soluble complexes [3] and another set of around 1,200 predicted complexes [4]. The size and complexity of the system does not easily lend itself to a readily readable overview of the entire system. Instead, a collection of resources are required to break the system into pieces that can be used for analysis and experimentation.

For example, WikiPathways is an open source pathway repository that is open to the community to create and modify pathway diagrams so that they can be shared with everyone in the community [5]. The WikiPathways database depicts biological processes and their connections to each other. The connections of elements within a pathway are shown as edges from one node to the next. These edges themselves have biological meaning that can be modelled and represented in WikiPathways [6].

For interoperability, WikiPathways also has a Resource Description Framework (RDF) set associated with it [7]. The RDF is the semantic representation of pathway diagram elements that are displayed and generated from the original Graphical Pathway Markup Language (GPML) in which WikiPathways stores the pathways. The WikiPathways RDF then includes both the graphical RDF (GPMLRDF) and the semantic elements of the RDF (WPRDF). The RDF allows users to go from creating an image of a biological pathway to trapping the elements and keeping them in a machine readable way and made available in a widely available way to be queried. The advantage of this is that it is also a linked data resource that can be queried by users at the WikiPathways SPARQL Protocol and RDF Query Language (SPARQL) endpoint, to query RDF databases (http://sparql.wikipathways.org/). This store of the WikiPathways RDF can be accessed both directly from the WikiPathways SPARQL endpoint, but also by remote requests via federated queries.

In order to represent connectivity between nodes in a pathway diagram, the meaning of a drawn line connecting nodes needs to be understood. WikiPathways RDF has connectivity information stored as point A is connected to point B. To a human looking at a pathway, it is more obvious what an arrow connecting two points means or what is implied by the arrow, but the RDF needs this stated explicitly if any inferences about how elements are connected is to be gleaned.

Furthermore, interactions can be either directed or undirected. Information about the direction and connectivity in a pathway diagram helps to explain the biological processes and therefore helps understand cause-effect relationships represented in the pathway. However, not all interactions have a clear direction: while the direction of a metabolic conversion follows chemical thermodynamics, interactions like the associations that exist in a complex are symmetrical and do not have a direction. Even more complex is a ligand binding, where the physical interaction is not only directed, but the interaction arrow also reflects the movement of the ligand. Therefore, it is important to know if an interaction has a directed route as part of a path and the RDF needs to preserve this information.

To ensure that pathway interaction drawings and notations can be biologically interpreted, the RDF needs to have standardized types for the interaction. That will allow users to query for all reactions of a similar (biological) type rather than worry about which notation was used in the drawing. WikiPathways supports several drawing notations, which can be general WikiPathways notations, MIM notations [8], and SBGN notations [9]. Based upon WikiPathways pathway ontology, these three can all be used and shown on WikiPathways. The available interactions themselves can be classified into nine different types: conversions, bindings, interactions, directed interactions, catalysis, transcription translation, complex bindings, inhibitions, and stimulations.

When interactions in various notations are normalized, more biological knowledge can be explored, and new questions answered. This interoperability effort makes it possible to gain implied knowledge from how a pathway diagram is drawn. For example, if two enzymes are catalyzing some chemical substrates in succession then there would typically not be a direct link or arrow drawn from one enzyme to another, but in order for the second enzyme to work the product from the first reaction must be present. This has the implication that the second enzyme is biologically downstream of the first enzyme, even though this interaction is not explicitly drawn. Having semantically clear directions and interaction types is essential to reach this conclusion from the RDF. Drawing of interactions with the WikiPathways and MIM notations can be done with the default installation of the PathVisio core [6], while SBGN needs a PathVisio plugin https://www.pathvisio.org/plugin/sbgn-plugin/. The PathVisio pathway editor thus makes it possible to annotate an interaction as a simple line with an arrowhead, as a MIM interaction, by default, or to create a SBGN drawing using plugins. It then becomes necessary to unify common types from the different graphical standards so that a MIM-Inhibition and a SBGN-Inhibition are represented as the same thing. After all, in both cases, the interaction is indicating an inhibitory effect of one entity upon another. Knowing the interaction types gives important context of the connection and the entities involved. A small note about how complexes are represented is also essential. All the entities are connected to each other with an undirected interaction. This keeps them all connected to each other as well as with any interaction that they are associated with as a complex.

The general interaction type is used to denote an interaction between data nodes and thus all interactions are of this type. A directed interaction, on the other hand, means there is a direction that says one data node is influencing another but the exact mechanism is not known or at least not described, by the pathway creator (author). Directed interaction is also the general data type for all interactions that have some directional information included. Therefore, all interactions have the type directed interaction except binding and complex binding, with the directed interaction itself being a child of the general interaction type. We therefore wanted to study to what extent we can derive knowledge from biological interactions, by semantically capturing biological meaning of interactions and harmonizing the notation in pathway drawings. We tested our hypothesis that this can be done by answering the following questions. First, can we translate graphically modelled biological knowledge to a semantic model of biological knowledge that harmonizes interaction types and captures implied directionality? And second, can we then take advantage of the semantic translation of the graphical biological knowledge to programmatically answer biological questions? For this latter question, we studied two specific biological questions as examples: in one example we look at MECP2 and explore alternative targets for this protein by looking for targets either upstream or downstream as they both have an effect on MECP2’s role. For the other example we studied how lipid metabolism is captured in the *Ganglio Sphingolipid Metabolism* pathway (wikipathways:WP1423, WikiPathways Project *et al.*, 2019).

MECP2 is a protein that is important in the methylation of DNA [10]. MECP2 has been shown to have an important role in lipid metabolism, without the MECP2 supporting this role defects have been shown in mice [11]. Mutations in the MECP2 gene have been linked to the development of Rett Syndrome [12]. This disease is responsible for a host of neurological developmental issues that affects infant development and is predominantly found in females [13]. The severity of the disorder is related to the specific mutation found in the individual patient [14]. Ehrhart *et al.* have already demonstrated the power of integrating different databases to retrieve links between genetic variants and phenotypes in the cases of a rare disease for which a limited amount of data is available in a single place [15]. Being able to look at alternative targets that are a part of the sequence of developments that lead to disorders such as Rett may end up helping us to expand the knowledge about alternative causes and treatment opportunities. The types of interactions described for MECP2 are a simple case of connectivity and directional information captured in WikiPathways and make a good example to demonstrate how this can be used to allow observation of upstream and downstream interactions.

The second example describes the metabolic regulation and modifications of Sphingolipids which are known to regulate several cell functions [16]. Sphingolipids are produced in the endoplasmic reticulum and the modifications of this lipid class alters the effect of the specific sphingolipid’s function [17]. The conversion of these metabolites from one form to another is regulated by enzymes that act as a catalyst for the reaction to take place. Sphingolipids also play a role in signal transduction [18]. The sphingolipids play an important role in the membrane of eukaryotic cells and are often associated with disorders in the degradation of the lipids [19]. This shows the importance of proper metabolite regulation and metabolism as disruptions can lead to serious diseases with high mortality rates. Understanding how these elements of the pathway are connected to one another and how they are directed helps to understand when the elements are not working correctly. There are also a large number of proteins that are known to interact directly with sphingolipids and are necessary for cell function [20]. In WikiPathways, these types of interactions are most often drawn with an arrow that shows the conversion of the metabolites from one form to another along with an associated catalysis reaction that is facilitated by an enzyme. Looking at how metabolism is modelled in wikipathways:WP1423 helps illustrate how these conversion and catalysis reactions are stored. Metabolism interactions are a more complicated set of interactions as an enzyme is typically seen acting on another interaction. The Sphingolipid metabolism pathway displays this more complex observation and allows the identification of the order of the enzymes found for potential upstream/downstream analysis.

## Methods

### WikiPathways Data

#### Interaction modeling

The interactions in WikiPathways are modeled by taking the graphical semantic information from the pathway diagram’s GPML representation. The harmonization of interactions is part of the WPRDF generation. This is done by analysis of the lines that represent interactions in the graphical representation, and using these to decide how the participants in the interactions are connected. All harmonized interactions have a unique ID, are linked to the participants, and have an interaction type as outlined in the introduction. If it is a directed interaction, it will also have a source and target node for the interaction. JUnit (https://junit.org/) was used to test the harmonization with several tests to verify that these connections in the GPML are being converted to RDF as expected. These tests include the original GPML and the expected outcomes as described in the code repository at https://github.com/BiGCAT-UM/WikiPathwaysInteractions/tree/master/FilesGPML.

#### Benchmark data

We used the RDF from the WikiPathways June 2019 release (https://zenodo.org/record/3369380). Both the WPRDF and the GPMLRDF components of the WikiPathways RDF were used in this study. To examine how pathways are drawn and used in WikiPathways, the analysis used only pathways from the Curated collection and only for *Homo sapiens*, and therefore excludes the Reactome collection [21, 22].

### Data Analysis

To aggregate and analyze the date, Jupyter notebooks running Python were used to collect all SPARQL queries that were used to query the WikiPathways SPARQL endpoint [23]. The notebooks are available from (https://github.com/BiGCAT-UM/WikiPathwaysInteractions/): *DataNodeStats.ipynb*, and *InteractionStats.ipynb*, and two for the two biological examples. The first two represent two different categories of queries. *DataNodeStats* retrieves information about data nodes in both parts of the WPRDF while the *InteractionStats.ipynb* file is used to return data about connectivity between the nodes in the WikiPathways RDF, representing both the semantic and the graphical RDF elements. *ExampleMECP2.ipynb* is the file for the query related specifically to the *MECP2 up and down stream targets* example. Finally, *ExampleLipidMetabolism.ipynb* is the notebook for the case of *Sphingolipid metabolism*. These notebooks and their use are here further described below.

#### Datanode Harmonization

Data nodes needed to be harmonized first in order to be able to examine the connections between the nodes. That allowed us to better estimate how well the interaction harmonization itself went. Therefore, we first looked at the data nodes. The *DataNodeStats.ipynb* notebook contains Python code to calculate a series of counts of data nodes, to estimate the amount of data and to get a baseline number of what we can expect for the success of conversion and harmonization of interactions. It is important to realize that for interactions where one of the participating data nodes is not in the WPRDF, then the conversion script will not to create the interaction due to the absence of participants. Therefore this interaction will not be found in the WPRDF and will affect our interaction counting. The notebook calculates the total number of data nodes of a certain type, in the Jupyter notebook section *Datanode Type Counts*, the number of GPMLRDF data nodes without a WPRDF datanode equivalent, the number and percentage of GPMLRDF data nodes of a specific type with a WPRDF data node equivalent, and the following corresponding numbers and percentages of GPMLRDF data nodes without a WPRDF data node equivalent. Furthermore, it determines the number of GPMLRDF data nodes of type Complex without WPRDF equivalents. This is used to specifically track which data nodes that are part of the complexes that can be found in the graphical elements part of the RDF but not found in the WPRDF, the biological component of the WikiPathways RDF, because the biological meaning of complexes is currently not always well-defined in pathway drawings in WikiPathways.

#### Interaction Harmonization

The *InteractionStats.ipynb* notebook contains code to calculate numbers that reflect the harmonization of interactions in the biological WPRDF, by taking into account the different drawing notations as a unified interaction type. The first few sections calculate overall statistics, the *Number of Non-Directed Interactions* (for example, bi-directional binding), *Count of Interaction Types* (reflecting the biological nature of the interaction), *Interaction Count with Unspecified Type*, and the percentage of non-directed interactions. The second set of sections characterize the nature of the interactions, e.g. *Interaction counts by participants*, *Participants for Interactions* (which reflects what datanode types are involved in an interaction), and *Identifier IDs by data source*.

In order to evaluate the conversion success, it calculates the complementary *GPMLRDF Interactions without a WPRDF equivalent* and *GPMLRDF Interactions with a WPRDF equivalent*, and the resulting percentages of success (see *GPMLRDF Interaction with Equivalent WPRDF out of Total GPMLRDF Interactions*). The GPMLRDF Interactions without a WPRDF equivalent was used to check to see how many interactions that are present in the graphical version of the RDF but not present in the biological WPRDF. The query for the percentage of WPRDF Interactions that are of unspecified type was used to see how accurately detailed the biological pathways are annotated. Finally, the percentage of non-directed interactions in the notebook calculated how many of the WikiPathways interactions are of non-directed type. When these are between metabolites and they may reflect missing biological annotation of directions.

### Usability

To test our hypothesis that we can harmonize the interaction information, we developed the Jupyter notebooks to first collect and query the data from the WikiPathways RDF. We then created several unit tests to validate how the modelled interactions behaved and to verify that they are created correctly. This ensures that when an interaction is drawn, we can keep track of the relevant semantic data represented, such as what nodes are connected to each other, what type of interaction is drawn between them, and how many nodes are expected to be part of the interaction. We can then test assumptions like interactions between metabolites should be directed conversions and interactions between different proteins should not be conversions and add aberrant results as curation tasks. We further tested with two biological examples if the harmonized semantified interactions give interpretable answers.

#### MECP2 up- and downstream targets

For the specific example used for MECP2 metabolism, the Jupyter notebook used a SPARQL query to the WPRDF. This query works by first searching for targets that are upstream or downstream of MECP2. The query then identifies data nodes that are associated with the HGNC symbol MECP2. The query in the Jupyter notebook finally finds associated pathways that have this HGNC symbol present and matches interactions that have MECP2 as a target in the interaction.

#### Sphingolipid metabolism

In the case of the specific example used for Sphingolipid metabolism, the Jupyter notebook used a SPARQL query to the WPRDF. The query retrieves the source portion of an interaction and displays its label. In the case of Sphingolipid metabolism, the queries identified enzymes that are associated with conversions in the pathway and returned results with the enzyme, interaction, the source metabolite and the target metabolite product.

## Results

To understand the amount of data that can be accessed via the RDF, we looked at the available RDF data for WikiPathways as GPMLRDF and WPRDF, the first being a direct translation of the original graphical depiction of the GPML files and the second covering the biological content. A quick count of the June 2019 release shows that the WPRDF used in this paper had 24,220 data nodes, and 13,928 interactions and is available at http://data.wikipathways.org. The subject of the paper is the interactions between data nodes, but we first need to understand that edges of a network connect datanodes to one another and so understanding the fundamentals of the biomolecular data nodes is necessary. This defines some context for the following results.

### Datanode Results

With regards to the data nodes, the most prevalent node type is the general Data-node type. It is the base type for any datanode, as described by the WikiPathways Vocabularies (https://vocabularies.wikipathways.org) and thus is used for every data node, it may include any of the descriptive data types. More specific but still generic, the GeneProduct type is the next most prevalent node type. These include explicitly typed proteins and RNAs and while the remaining GeneProduct typed nodes are not specified further. Table 1 illustrates the size of the WikiPathways semantic RDF part and the types of nodes present in WikiPathways. There are a total of 24,220 data nodes, the majority of which are gene products. Proteins are the next common type followed by metabolites and RNA. There are also Complex nodes to represent clustered groups of other node types, specifically proteins, gene products, and RNA. Pathways are not typed as Datanode in the WPRDF, which is why the value is blank in the table. Overall, 5.3% of GPMLRDF data nodes do not have a WPRDF data node equivalent and thus 94.7% of the GPML data nodes are found in both parts of the RDF.

**Table 1.** Datanode Type Counts, as defined by the WikiPathways ontology. The Datanode counts for each type of node.

|  | Datanode Type | Count (WPRDF) | Count (GPMLRDF but not WPRDF) |
| --- | --- | --- | --- |
| 0 | Datanode | 24220 | — |
| 1 | GeneProduct | 18832 | 417 |
| 2 | Protein | 6881 | 165 |
| 3 | Metabolite | 3547 | 223 |
| 4 | RNA | 1106 | 48 |
| 5 | Complex | 33 | 18 |
| 6 | Unknown | — | 274 |
| 7 | Pathway | — | 224 |

Also seen in Table 1 are the data nodes that are found in the GPMLRDF but not found in the WPRDF. The reason typically is that the node exists but is not linked to a clear biomolecular database identifier, in other words we do not know exactly what it is. Datanodes are any node type in the pathway diagram and the count of gene products also includes proteins and RNAs as these are specifications of the products produced. Complexes are a combination of several other node types that form a unit with one another. We can also see how many data nodes are found in both parts of the RDF.

If we specifically look at some examples of data nodes that are present in the GPMLRDF but not carried over to WPRDF, we can see a list of eighteen complex data nodes, and the details of these are given in Table 2. This second table also includes the labels for the complexes, shedding some light on which complexes were not transferred over to the semantic portion (WPRDF) of the RDF from the graphical portion (GPMLRDF). There is no clear pattern with one reason for why the complexes are converted in some cases but not others.

**Table 2.** The Complex GPMLRDF datanodes without WPRDF equivalents. This is the same 18 complexes identified in Table 1.

|  | Complex Datanodes | Complex Labels |
| --- | --- | --- |
| 0 | WP2806_r97328/Datanode/a680a | C1q |
| 1 | WP2806_r97328/Datanode/b45a6 | C3bB3bP |
| 2 | WP2806_r97328/Datanode/e4901 | C3bB3b |
| 3 | WP2806_r97328/Datanode/f51e9 | C4b2b |
| 4 | WP2806_r97328/Datanode/f7d7c | C3bBbP |
| 5 | WP2806_r97328/Datanode/fd0cb | C3bBb |
| 6 | WP2806_r97328/Datanode/fd947 | C4b2b3b |
| 7 | WP2509_r92315/Datanode/c040f | mTORC1 |
| 8 | WP391_r71373/Datanode/fc7a3 | PKA |
| 9 | WP4197_r95608/Datanode/d7ca2 | DRIP150 |
| 10 | WP2583_r97638/Datanode/a8ece | B7-1/ B7-2 |
| 11 | WP2583_r97638/Datanode/c951f | PD-L1 |
| 12 | WP2583_r97638/Datanode/da6cd | MHC peptide |
| 13 | WP2583_r97638/Datanode/f7b20 | B7-1/ B7-2 |
| 14 | WP2586_r91687/Datanode/b328d | DRE region |
| 15 | WP2586_r91687/Datanode/c3d69 | TATA |
| 16 | WP3601_r89202/Datanode/c8eb7 | Chylomicron |
| 17 | WP3601_r89202/Datanode/e3358 | IDL |

When we do this evaluation for the pathways of the two use cases, we find that for wikipathways:WP4312, which pertains to MECP2, there is 1 gene product type data node that is found in the GPMLRDF but not found in the WPRDF. This represents 1 gene product out of 148 other gene products that were found in the WPRDF and out of 152 total data nodes found in the WPRDF. In the instance for wikipathways:WP1432, which is related to sphingolipid metabolism, there is 1 metabolite that is found in the GPMLRDF but is not found in the WPRDF. This is 1 metabolite from 38 total metabolites found in the sphingolipid metabolism pathway and out of 62 data nodes found in the WPRDF for wikipathways:WP1432.

In the Additional file 1 there are tables with examples of data node types that are found in the GPMLRDF but not in the WPRDF for various pathways (as counted in Table 1). In this file, the top ten results are retrieved along with the table to give some idea why they may not be translated. In Additional files 2 and 3 there are tables for the data node counts for the specific WikiPathways example pathways of MECP2 and Sphingolipid metabolism.

### Interaction Results

Similar to what we did for the data nodes, we calculated non-directed interactions and non-specific interactions along with the specific interaction types and counts. Non-directed interactions being all interactions that do not have any directional information, such as in the case of a binding event. Non-specific, on the other hand, means that an interaction does not even have a specified non-directed interaction like a binding.

First, we identified nine interaction types. The overview of mappings to WPRDF of the GPML interaction types that can be found in WikiPathways, is available from https://github.com/BiGCaT-UM/WikiPathwaysInteractions/tree/master/FilesGPML. This GitHub repository contains example GPML files for each interaction type that can be found at https://vocabularies.wikipathways.org/, along with an example of what the interactions look like in GPML, as well as files with statistics about the interaction as it appears in the WPRDF. These numbers are used in the JUnit tests to verify that the different models are harmonized into the single interaction model in WPRDF. These tests are now available as part of the regular testing of RDF generation (see https://github.com/wikipathways/GPML2RDF, *src/test/java/org/wikipathways/wp2rdf/interactionTests* folder). When we look at the full WPRDF, the types of generic non-directed and nonspecific interactions can be seen. Out of a total of 13,928 interactions, 3,299 (23.7%) were non-directed and 2,631 (18.9%) were non-specific (see Table 3). Thus 7,998 (57.4%) of the interactions have some sort of direction information. The number of non-specific interactions can be either an indication that there is just not sufficient evidence to explain what the interactions are or that better curation is necessary. Examples of how interactions are drawn in WikiPathways can be seen in Figure 1.

**Table 3.** Interaction Type Counts, as defined in the WikiPathways ontology.

|  | Interaction Type | Count (WPRDF) | Count (GPMLRDF but not WPRDF) |
| --- | --- | --- | --- |
| 0 | Interaction | 13928 | — |
| 1 | DirectedInteraction | 10629 | — |
| 2 | Conversion | 1079 | — |
| 3 | Inhibition | 1021 | — |
| 4 | Catalysis | 1000 | — |
| 5 | ComplexBinding | 668 | — |
| 6 | Binding | 668 | — |
| 7 | Stimulation | 661 | — |
| 8 | TranscriptionTranslation | 232 | — |
| 9 | NonDirected | 3299 | — |
| 10 | NonSpecified | 2631 | — |
| 11 | Unknown | — | 11081 |

**Figure 1.**
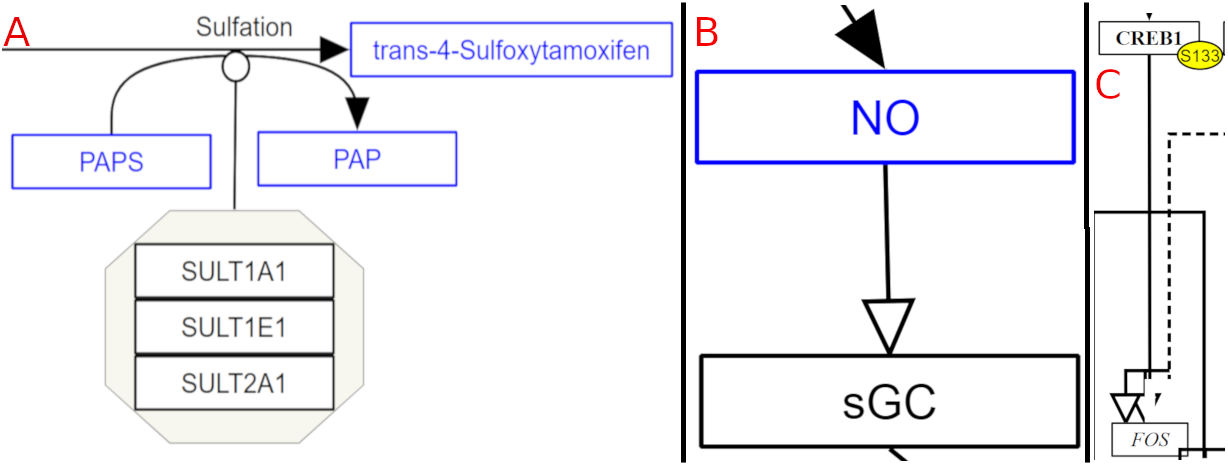
Interaction types that are not found in the specific examples below. A shows a complex binding found in Tamoxifen Metabolism (wikipathways:WP691). In B a stimulation interaction is shown for Endothelin Pathways (wikipathways:WP2197). C shows transcription translation interaction in Corticotropin-releasing hormone signaling pathway (wikipathways:WP2355).

Only a small percentage of the interactions have associated identifiers. Having such identifiers can make it easier to find information about the provenance of that interaction occurring in a pathway and it is useful for linking experimental data or modelling results to the pathway or to find descriptions of the interactions in external resources. Table 4 contains provenance information about the databases to which identifiers for interactions refer. UniProt-TrEMBL has the most interactions represented in WikiPathways. There were some unexpected database links. Sources like *kegg.compound* and *ChEBI* are not expected to have interaction data information but are included because the user identified them as the database resource for the interaction. These unexpected sources come from two pathways, wikipathways:WP3634, and wikipathways:WP3535. Currently, most interactions do not have any database identifier associated with them, the main reason is probably that the mechanism to add these is relatively new.

**Table 4.** Interaction Identifier ID counts by data source.

|  | Database Source | Interactions |
| --- | --- | --- |
| 0 | Uniprot-TrEMBL | 213 |
| 1 | Other | 71 |
| 2 | KEGG Pathway | 18 |
| 3 | pato | 8 |
| 4 | kegg.compound | 8 |
| 5 | ChEBI | 6 |
| 6 | KEGG Reaction | 3 |
| 7 | WikiPathways | 2 |
| 8 | XMetDB | 2 |
| 9 | SPIKE | 2 |
| 10 | BIND | 1 |

Finally, to further characterize the interactions present, Tables 5 and 6 provide examples of the makeup of the interactions seen in WikiPathways. Table 5 shows example Interaction IDs, along with their interaction types, and what type of Data-node type is participating in the interaction. And Table 6 shows the profile with the interaction participants and a count of how many times this interaction profile was counted in WikiPathways and the type of these interactions.

**Table 5.**
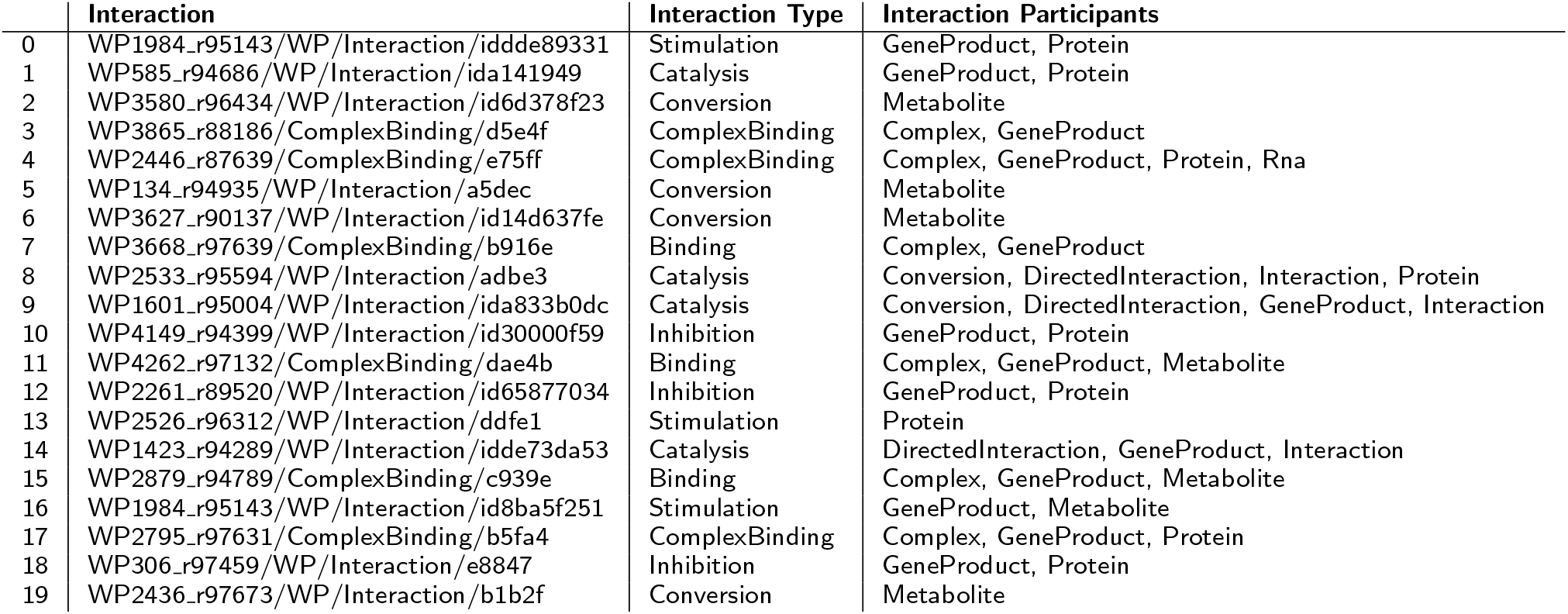
Participants for Interactions. Twenty example interaction syntaxes shown in table below.

**Table 6.** Top 20 most occurring directional interactions by participants combination. The most abundant interaction is a directed interaction between two metabolites.

|  | Interaction Participants | Count | Type |
| --- | --- | --- | --- |
| 0 | Metabolite, Metabolite | 2367 | DirectedInteraction |
| 1 | GeneProduct, GeneProduct | 1436 | DirectedInteraction |
| 2 | GeneProduct, Protein, GeneProduct, Protein | 1230 | DirectedInteraction |
| 3 | Metabolite, Metabolite | 873 | Conversion |
| 4 | Metabolite | 420 | DirectedInteraction |
| 5 | GeneProduct, Protein, GeneProduct | 389 | DirectedInteraction |
| 6 | GeneProduct, Protein | 374 | DirectedInteraction |
| 7 | GeneProduct, GeneProduct, Protein | 329 | DirectedInteraction |
| 8 | GeneProduct | 300 | DirectedInteraction |
| 9 | GeneProduct, Protein, Protein | 292 | DirectedInteraction |
| 10 | DirectedInteraction, Interaction, GeneProduct | 275 | DirectedInteraction |
| 11 | GeneProduct, GeneProduct | 272 | Inhibition |
| 12 | Protein, Protein | 262 | DirectedInteraction |
| 13 | Metabolite, GeneProduct | 246 | DirectedInteraction |
| 14 | DirectedInteraction, Interaction, GeneProduct | 240 | Catalysis |
| 15 | GeneProduct, Metabolite | 213 | DirectedInteraction |
| 16 | GeneProduct, DirectedInteraction, Interaction | 151 | DirectedInteraction |
| 17 | Protein, Protein | 142 | Stimulation |
| 18 | DirectedInteraction, Interaction, Conversion, GeneProduct | 140 | Catalysis |
| 19 | DirectedInteraction, Interaction, Conversion, GeneProduct | 140 | DirectedInteraction |

When this notebook was applied to the pathways of the two use cases, we found that for wikipathways:WP4312, which pertains to MECP2, there are 5 interactions that are found in the graphical GPMLRDF but not found in the semantic WPRDF. This represents 5 interactions out of 45 non-specified interactions that were found in the WPRDF and out of 37 directed interactions found in the WPRDF. In the instance for wikipathways:WP1432, which is related to sphingolipid metabolism, there are 24 interactions that are found in the GPMLRDF but is not found in the WPRDF. These are 24 interactions found in the GPMLRDF while 49 directed interactions were found in the WPRDF for the sphingolipid metabolism pathway, wikipathways:WP1432.

In the Additional files 2 and 3 tables can be found for the interaction counts of the two specific pathways for MECP2 and Sphingolipid metabolism. These contain the types of interactions found in these pathways as well as how many interactions were found in the GPMLRDF but not in the WPRDF resources for WikiPathways as described above.

### MECP2 up and down stream targets

We created Jupyter notebooks to evaluate the example pathways, as described in the Methods section. The SPARQL queries used in the Jupyter Notebooks will return the interactions that have MECP2 as a participant and then the associated upstream source of the interaction or the associated downstream target of MECP2 and can be found in Table 7. Figure 2 shows examples of the directed nature of influences by MECP2. The query identified ten gene products that are known to influence or be influenced by MECP2. Three gene products were upstream of MECP2 and have an influence on MECP2, while the other 7 gene products were downstream of MECP2 and indicate that they are influenced by MECP2. This basically captures the semantics of the biological meaning of the pathway, a rare disease caused by a damaged gene that has a variety of effects and interactions.

**Table 7.**
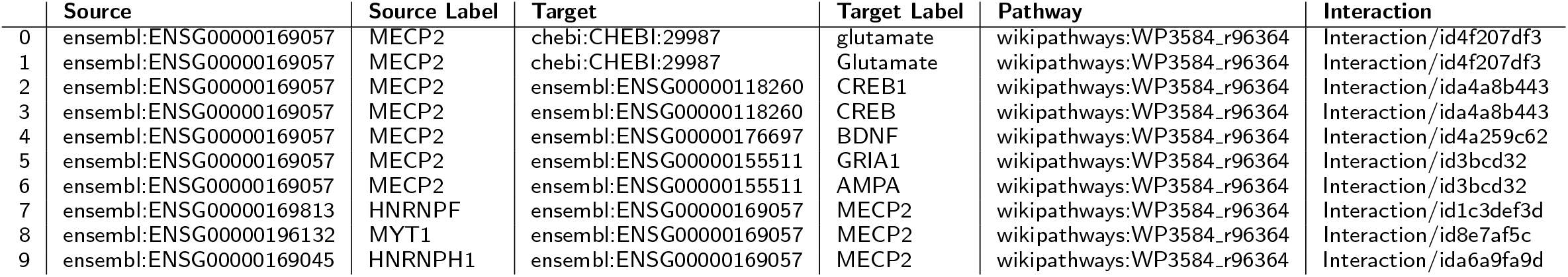
MECP2 Upstream and downstream targets.

**Figure 2.**
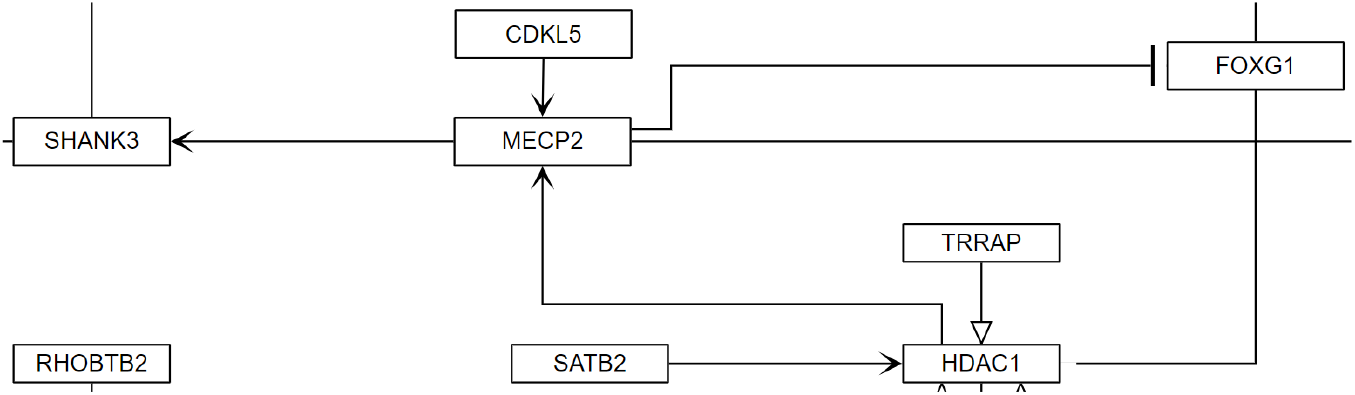
Example of direct interactions of gene products that both influence MECP2 and are influenced by MECP2 from Rett syndrome causing genes (wikipathways:WP4312). In this example, MECP2 is being influenced by HDAC1 and CDKL5. MECP2 then in turns influences SHANK3 and inhibits the activity of FOXG1.

### Sphingolipid metabolism

For Sphingolipid metabolism, a Python script was devised that queries the WPRDF for WikiPathways pathway wikipathways:WP1423, Ganglio Sphingolipid Metabolism, and returns a table with directed interactions that have an enzyme that is catalyzing the reaction. The query limits results to wikipathways:WP1423 as a matching criteria, then finds interactions that are annotated as being a catalysis reaction. It retrieves the associated protein for the catalysis along with the interaction that is being acted upon. Finally, the query also retrieves the source (substrate) and target (product) for the directed interaction that was being catalyzed. Figure 3 shows an example enzymatic reason. The results of the query are shown in Table 8, five conversion annotated interactions in this pathway were returned.

**Table 8.**
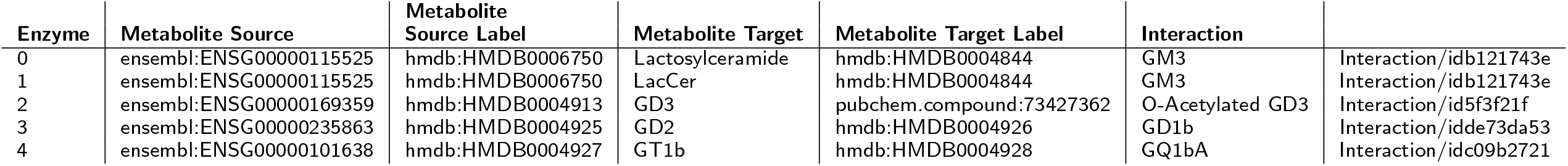
Sphingolipid Conversion Interactions.

**Figure 3.**
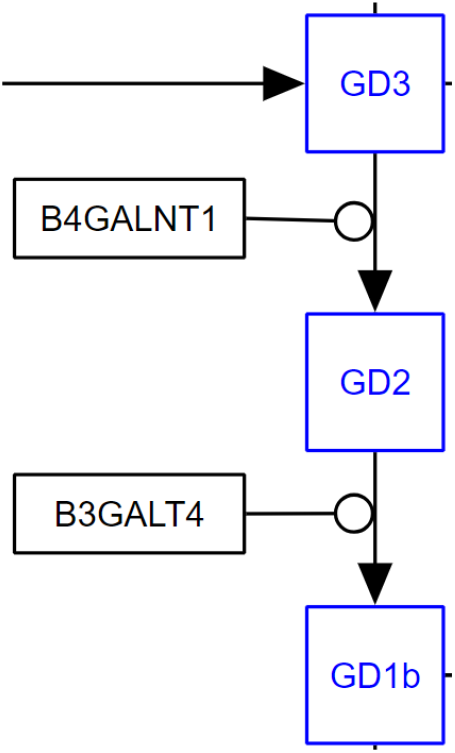
Representation of conversion of different sphingholipids to their products and the relevant enzyme catalyzing the reaction from the Ganglio Sphingolipid Metabolism pathways (wikipathways:WP1423). In this case, GD3 is converted to GD2 by the enzyme B4GALNT1. GD2 is then in turn converted to GD1b and catalyzed by B3GALT4.

## Discussion

The analysis in this paper only involves human pathways on WikiPathways from the original, non-Reactome, collection. For other species, the results would have been affected by the more limited curation effort that has been spent on those in general. To allow us to do meaningful interaction analysis we need to have sufficient information about the interactions and their participants. Generally, a data node might be found in the graphical portion of the RDF and not in the semantic portion because of incorrect annotations, because the curator really meant to add something atypical, like an organ, or because of a failure by the conversion scripts to successfully convert the graphical information into semantic information.

Interaction types were harmonized by the scripts to turn pathway graphical information into semantic data if there was an appropriate analogue and drawing for the different notation types. This allows for example, a user to draw either a SBGN, a MIM, or a general WikiPathways inhibition drawing to all have a harmonized interaction type called wp:Inhibition. In this example, since all three different notation types have the same biological meaning of indicating an inhibition event, it allows the user the flexibility to draw the pathway in the notation they are most comfortable using and still preserving the meaning of the interaction edge.

For the more curated human pathways, we find that gene products that are in the GPMLRDF but not in the WPRDF, typically these are nodes that do not have a selected database resource type, like Ensembl or NCBI Gene. From Table 1 we learn that most of the data nodes already do have enough information to be included in the semantic part of the RDF. Future curation tasks to identify appropriate sources for the data nodes with missing annotations would enable them to become part of the semantic information.

Three further examples of existing problems with data nodes exist for nodes of unknown type, pathway nodes, and complex data nodes. The unknown nodes do not have an associated data type or an associated database. Pathways nodes are currently not part of the WPRDF data model.

In the case of data nodes for complexes, there were only 18 complex nodes that do not have an equivalent in the semantic information. These complex data nodes also share the problem of missing database resource and data node identifiers.

We also saw how data node types and interaction types complement each other. For example, Table 4 shows specific interactions as well as the type of the interaction and the interaction’s participants. This can also be a useful aid in helping to identify areas of curation that need to be addressed. For example, if the participants retrieved for a conversion reaction are metabolites then this makes sense, but if the participants are proteins then there is a possibility that a post-translational modification is described put is also possible that the user used the wrong annotation for the interaction type, especially when the two proteins are known to be derived from different genes. Based upon the results summarized in Table 5, we can get an estimate of what combinations of participant and interactions types are most prevalent. This gives us an indication of the accuracy of the data. For example, we found a large number of directed interactions connect two metabolites without specific type These likely are conversions but still miss that typing.

We further found that one reason why interactions are captured by the GPML-RDF but not the WPRDF is because some interactions are lines connecting one or more text labels. These are not converted into the semantic layer. The WikiPathways database also allows information added as graphical annotation for the user to better understand a pathway diagram and to provide background information. This type of graphical annotation is not incompletely curated data and is not meant to show up in the WPRDF.

A third reason why some interactions are not captured in the semantic layer is because one of participants is a user defined group or complex. Ideally, when the participant really is a complex, then that complex itself should be identified with an external identifier like one from the Complex Portal at EBI (https://www.ebi.ac.uk/complexportal/home) [24]. In that case it is clear that all elements of such a complex are involved in the reaction, although the curator may still have made clear that one element is directly involved. In that case, the interaction will be graphically connected with an element inside the complex.

Also in the GitHub repository is a directory titled *pastReleases* with tables of values for the queries that were performed on the November 2016 release of the WikiPathways RDF as a comparison to the June 2019 release used in this paper. Additional file 4 is also included as a zip file for the results of the June 2016 release. What is reflected in this comparison is that there is ongoing growth of the WikiPathways database and its semantic descriptions which sees a 43.8% increase in datanodes and an 23.3% increase in interactions from the 2016 release to the more recent release. All datanode types and interactions saw an increase in the later release compared to the earlier release, except for the case of stimulation interactions. This value went down between the releases as a result of curation efforts that identified that several of the interactions annotated as stimulations were incorrectly typed as such. Because of this curation the interactions were re-typed as their appropriate interaction type and thus we see a decrease in their number of interactions.

There is an ongoing discussion on user defined groups too, e.g. on how those should be connected and represented in the RDF as there might not be a single solution to address all the use cases of user groups. For example, these user groups often represent a class of enzymes that are all capable of catalyzing the same reaction, this can be seen in the example of the Sphingolipid Metabolism pathway, wikipath-ways:WP1423. Several intended interactions are not included in the WPRDF since the participants belong to a group of isoenzymes and will not be found in SPARQL query results. For this case, a simple solution would be to connect each element of the group via a duplicate interaction that is annotated as a catalysis towards the conversion, but not connect the isoenzymes to each other as is implied in the case of a biological complex. However, a user group could currently be any sort of convenient grouping and so this solution would not be a catch all solution for all groups, and further specifications would have to be included in the WikiPathways drawing options set itself.

The modelled biological knowledge of WikiPathways has previously been reported in the Waagmeester *et al.* paper [7]. During that analysis, the first release of WPRDF was explored to determine how elements were connected to one another in that semantic part of the RDF. As discussed above, there were many interactions that are drawn in the pathway and in the graphical information about a pathway but not found in the semantic layer. This was partly addressed by curation efforts that made sure that data nodes are drawn, typed and identified correctly and interactions are drawn for instance from anchors of the data nodes to another anchor in the drawing program. Overall 56% of interactions in the graphical information is now represented in the semantic portion.

Nevertheless, as was shown in the two biological examples above, it is possible to take advantage of the semantic information in the RDF to answer relevant questions. For MECP2, known to be a core epigenetic regulator, it was possible to identify MECP2 in pathway diagrams and then use connectivity information to find which other elements have a direct influence upon it and which elements MECP2 influences directly. In Sphingolipid metabolism, conversion of metabolites from one form to another by a catalysis reaction were shown. This has interesting implications as it is then possible to expand this knowledge to infer information about the hierarchy of enzymes in this pathway. Meaning that, for example, GD3 is converted to GD2 by enzyme B4GALNT1 and GD2 is converted to GD1B by enzyme B3GALT4. This means that anything that acts upon and affects the activity of the upstream B4GALNT1 enzyme, will also affect the conversion of GD2 to GD1B by B3GALT4 through influence on substrate availability. This is more of an indirect influence of one element upon another but it is possible to then retrieve these indirect interactions.

## Conclusion

It was demonstrated that most of the graphical biological knowledge from WikiPathways is modelled in the semantic layer (WPRDF) of WikiPathways RDF with the semantic information intact and connectivity information preserved. This semantic translation allows us to answer biological questions. The MECP2 example shows directional regulatory information captured by the WPRDF, and for the other example of Sphingolipid metabolism complex successive biochemical reactions are captured. MECP2 involvement in regulatory, epigenetic interactions has implications for the understanding of the rare disease Rett syndrome. Sphingolipids are important parts of cell function and structure. Being able to evaluate the order in which biological elements affect each other allows, for example, the identification of up or downstream targets that will have a similar effect when modified.

The usability of the WikiPathways pathway and connectivity information has shown to be useful and has been integrated into platforms such as the Open PHACTS Drug Discovery Platform [25]. Improvements in WikiPathways curation and in the conversion to WikiPathways RDF support these other platforms and will allow giving a more complete picture of connectivity in biological systems. Continued curation efforts will incrementally improve many of the shortcomings of data and will continually make the semantic information better. Efforts to improve on the conversion scripts can address lost connectivity information that is for instance the result of using groups and complexes. Pathways themselves are also continually being added to WikiPathways and will continue to add to the richness of knowledge of biological interactions.

## List of Abbreviations

RDF: Resource Description Framework
GPML: Graphical Pathway Markup Language
GPMLRDF: RDF for Graphical Pathway Markup Language
MIM: Molecular Interaction Map
SBGN: Systems Biology Graphical Notation
WP: WikiPathways
WikiPathways RDF: The combination of GPMLRDF and WPRDF
WPRDF: RDF for WikiPathways
SPARQL: SPARQL Protocol and RDF Query Language
KEGG: Kyoto Encyclopedia of Genes and Genomes
HGNC: HUGO Gene Nomenclature Committee

## Ethics approval and consent to participate

Not Applicable

## Consent for Publication

Not Applicable

## Availability of data and Material

The Jupyter Notebooks used for the analysis can be found at https://github.com/BiGCAT-UM/WikiPathwaysInteractions/tree/master.

The RDF can be found as a data dump at http://data.wikipathways.org/current/rdf/.

The WikiPathways SPARQL endpoint is found at http://sparql.wikipathways.org/.

The Zenodo archive of the WikiPathways data is found at https://zenodo.org/record/3369380 [26].

## Author’s contributions

RM developed the Jupyter notebooks, SPARQL queries, and GPML interaction examples, analyzed the results, and designed the study. EW and CE supervised the project and contributed to the writing. AB, MK, and AW were involved in the design of the interactions. All authors read and approved the final manuscript.

## Acknowledgements

A thank you to the authors and curation teams at WikiPathways, they require a special acknowledgement for their maintenance of the primary resource used for analysis. The work done here was also inspired by the work done in collaboration with the Open PHACTS Discovery Platform.

## Additional Files

Additional file 1 — NotFoundInWPRDF.csv

Table for the top 20 datanodes that are found in the GPMLRDF but not in the WPRDF presented in the CSV file format accessible through most spreadsheet programs.

Additional file 2 — MECP2Stats.csv

Table for the conversion statistics of the MECP2 pathway wikipathways:WP4312 in the CSV file format accessible through most spreadsheet programs.

Additional file 3 — SphingolipidStats.csv

Table for the conversion statistics of the Sphingolipid Metabolism pathway wikipathways:WP1423 in the CSV file format accessible through most spreadsheet programs.

Additional file 4 — 201611RDFResults.zip

Zip file for the tables of values for the queries that were performed on the November 2016 release of the WikiPathways RDF.

